# Mating allows asexual female wasps to produce offspring both sexually and asexually, but their sexual daughters have reduced fecundity

**DOI:** 10.64898/2026.09.16.751999

**Authors:** R.A Boulton, M. Raj, J Anderson, D. J Parker

## Abstract

Sexual reproduction is widespread despite being less efficient than asexual reproduction. While its long-term evolutionary benefits are well recognized, it remains unclear why facultative sex, which should allow lineages to balance the advantages of both sexual and asexual reproduction, is not more common. One potential reason is that intermittent sex imposes novel costs compared to obligate sex, limiting the persistence of facultatively sexual lineages and constraining introgression. We test this possibility using the sexually polymorphic aphid parasitoid wasp *Lysiphlebus fabarum*. We reared offspring from females who reproduced asexually or by facultative sex for three generations (without sex after the founding generation). We genotyped F1 females to confirm facultative sex and used antennomere number, a phenotypic marker associated with reproductive mode, to test whether mixed sexual-asexual genotypes could be tracked across generations. We found multiple barriers to facultative sex. Mating did not always result in offspring with sexual-asexual genotypes, and when sex did occur, broods were usually comprised of asexually and sexually produced offspring. Additionally, lineages founded by females produced through facultative sex showed reduced fecundity compared to asexually produced broods. We saw declining frequencies of the sexual antenna phenotype in the generations after facultative sex, suggesting that genotypes generated by facultative sex are selected against. Overall, our results provide empirical support for the idea that facultative sex can impose costs that persist beyond the initial sexual event. These costs, together with low realised rates of sex, may limit long-term introgression and explain why facultative sex appears rare compared to obligate sex.

## Introduction

Sexual reproduction dominates eukaryotic life despite substantial costs, most notably the “twofold cost of males” (Maynard-Smith 1978): if a sexual female produces ∼50% sons, she has half the reproductive output of an equivalent asexual female producing only daughters. Given this demographic disadvantage, asexuals should dominate,yet sexual reproduction remains widespread in nature (Lehtonen et al. 2012).

Sex must therefore confer substantial benefits to explain its ubiquity (Neiman & Schwander 2011). Theory has proposed many such benefits, which can be broadly separated into mutational models, where sex purges deleterious mutations and accelerates the spread of beneficial alleles (Hartfield & Keightley 2012; Kondrashov 1994), and environmental models, where recombination and genetic diversity improve performance under changing or antagonistic selective conditions (Lively & Morran 2014; Neiman et al. 2018).

The genomics era has provided powerful tools to better understand the evolution of sex and has added nuance to the question, showing that reproductive mode often lies on continuum rather than forming discrete sexual–asexual categories. We now know that many species previously thought to be obligately asexual engage in occasional or facultative sex (Hartfield & Keightley 2012; Hartfield 2016; Kokko 2020; Frietas et al. 2023; Molinier et al. 2025; Parée & Teotónio 2025; Chung et al. 2026). Notably, in species with polymorphic reproductive modes, cryptic facultative sex opens up the possibility of gene flow between obligately sexual and asexual populations, meaning that asexuals can access the long-term benefits of sex without consistently paying the short-term demographic costs (Kokko 2020).

At face value, facultative sex appears to be the ‘best of both worlds’, combining the genetic benefits of sex with the demographic efficiency of asexual reproduction. This raises the question of why facultative sex is not more widespread than obligate sex (Neiman et al. 2014; Burke & Bonduriansky 2017; Kokko 2020; Wilner et al. 2025). One explanation is that the immediate costs paid by sexual individuals may be greater under facultative than obligate sex (Lehtonen et al. 2012; Parée & Teotónio 2025). These costs arise from processes inherent to sexual reproduction (such as the disruption of coadapted gene complexes, changes in heterozygosity, the generation of incompatibilities, and sexual conflict), but their effects may be amplified or more likely to arise when sex is intermittent (Lehtonen et al. 2012; Hartfield 2016; Burke & Bonduriansky 2017; Parée & Teotónio 2025).

To understand the evolutionary significance of facultative sex, it is necessary to determine how these costs translate into longer-term outcomes. In particular, the extent to which sexually produced individuals in predominantly asexual populations persist and contribute to gene flow, or whether they are selectively disadvantaged and fail to compete with established asexual lineages remains unclear. If the latter is true, then the potential for facultative sex to facilitate meaningful introgression may be limited because hybrid genotypes fail to persist. In this case, longer-term consequences of facultative sex, whether beneficial, neutral, or deleterious, may rarely be realised, as recombinant lineages are lost before they can contribute to evolutionary change.

Addressing this question is challenging in natural populations, where facultative sex is often rare or cryptic and can be hard to detect (Pieszko et al. 2025). As a result, it can be difficult to disentangle whether occasional sex leads to sustained introgression or simply produces transient, low-fitness genotypes that are rapidly lost. To resolve this, experimentally tractable systems that can be followed across multiple generations are needed. By combining controlled crosses with multi-generational fitness assays and genotyping, it is possible to assess the success or failure of sexually produced lineages, and to quantify the extent to which recombined genotypes persist following facultative sex.

We used such a tractable experimental system, the sexually polymorphic aphid parasitoid wasp *Lysiphlebus fabarum*, to follow the fate of lineages produced under facultative sex over multiple generations. Previous work has shown that facultative sex may carry greater short-term costs than obligate sex or asexual thelytoky in this system (Boulton 2025) and so we expect lineages produced under facultative sex to be less successful. Our primary aim was to test whether facultative sex generates lineages that persist and contribute to introgression, or whether they are lost over time due to reduced fitness. To do this, we induced facultative sex in normally asexual (thelytokous) females by mating them with males produced by sexual (arrhenotokous) females, with matched unmated controls from the same asexual lineages. Unmated F1 daughters were then used to propagate the same families for three generations. F1 offspring were genotyped to confirm facultative sex in a subset of families, and antennomere number (see Fig. 1 and Table 1) was assessed as a potential phenotypic marker for tracking the persistence of sexual alleles through predominantly asexual populations. This experimental design allowed us to test whether (i) lineages produced through facultative sex have lower fitness and are more likely to go extinct than purely thelytokous lineages, and (ii) whether any signal of sexual reproduction persists across generations in surviving lineages.

**Table 1.** Origin and characteristics of the four lines used in this study.

| Line | Reproductive mode | Collection location | Year | Notes |
| --- | --- | --- | --- | --- |
| IL09-348 | Asexual (thelytokous) | Geneva, Switzerland | 2009 |  |
| IL09-402 | Asexual (thelytokous) | Langenthal, Switzerland | 2009 | Notable for near-absence of 12-segment females |
| MP24-0204 | Asexual (thelytokous) | Manchester, UK | 2024 |  |
| Sexual population | Sexual (arrehenotokous) | Mixed sites across Switzerland | 2012 | Large, genetically diverse outbred population |

**Figure 1.**
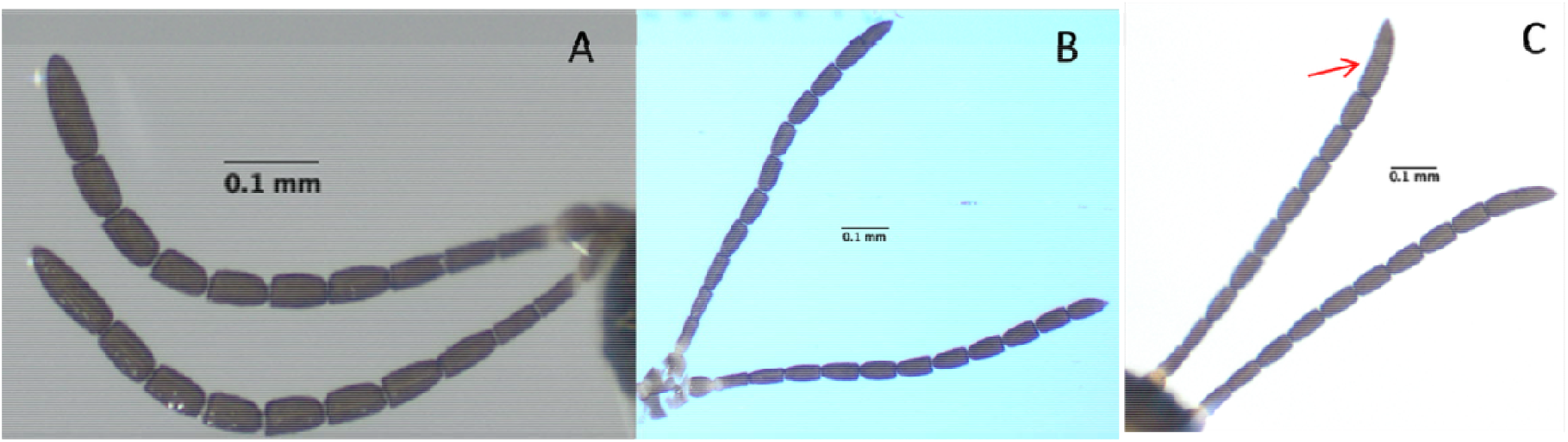
**Lysiphlebus fabarum** antenna types (A) Sexual-12; (B) Asexual-13 and (C) Mixed. In C, the arrow shows where the apical antennomere has only partially split (intermediate between 12 and 13 segments) while the other antenna is of the sexual-12 type. Note that the scape and pedicel which make up the first segments are counted as separate antennomeres.

## Methods

### 2.1 Study system

*Lysiphlebus fabarum* is a solitary koinobiont parasitoid that attacks several aphid species, with a preference for early-instar *Aphis fabae* (black bean aphid) nymphs (Najafpour et al. 2016). The species contains both obligately sexual (arrhenotokous) and thelytokous asexual lineages, sometimes occurring in sympatry (Belshaw et al. 1999; Sandrock et al. 2011). Asexuality is caused by central-fusion automixis which is linked to a recessive allele at a single-locus region (*Lysi07*), likely originating ∼0.5 MYA (Belshaw et al. 1999; Sandrock & Vorburger 2011). A microsatellite marker, *Lysi07*, can be used to distinguish sexual from asexual genomic backgrounds (Sandrock et al. 2007). Although asexuality is generally obligate, low-frequency facultative sex has been suggested (Belshaw et al. 1999; Sandrock & Vorburger 2011) and recently confirmed in the lab (Boulton 2025). These rare sexual events can be detected genetically through heterozygosity at *Lysi07* or through mixed multilocus genotypes (Boulton 2025).

### 2.2 Stock phenotyping and antennomere categories

While screening material from a previous experiment, we noticed differences in antennomere number between sexual and asexual lines: sexual females typically have 12 antennomeres per antenna and asexual females commonly have 13. In addition, many females did not fit clearly into either category. Some females had one 12-segment and one 13-segment antenna, and others had split terminal segments where the distal antennomere was only partially divided (Fig. 1). All sexually produced males we looked at had 14 antennomeres. Occasionally, asexual females will produce a male (Sandrock & Vorburger, 2011), we have only recovered 2 of these in the lab and both had 15 antennomeres.

To quantify variation in antennae, we phenotyped females from stock populations of the four lines used in this study: 83 females from three asexual lines and 199 females from an outbred sexual population (see Table 1). Each female was scored as:

□ **Sexual-12**: both antennae with 12 antennomeres (Fig. 1A)
□ **Asexual-13**: both antennae with 13 antennomeres (Fig. 1B)
□ **Mixed**: anything in between (12/13 combination or split terminal segments; Fig. 1C)

Mixed types appeared in all lines (∼10–25%) but were not clearly related to reproductive mode. Because the sexual-12 and asexual-13 types showed clear differences between sexual and asexual populations (see 3.1 results), the main analyses focus on these two categories.

### 2.3 Laboratory rearing

All rearing took place in the University of Stirling Controlled Environment Facility under 20°C, 70% RH, 16:8 L:D. The sexual population was maintained in large BugDorm™-4F3030 cages (∼200 adults per generation) on **A. fabae** feeding on broad beans (**Vicia faba)**. Asexual lines and all experimental families were kept in modified “culture cups”: a plastic cup with a hole at the base inserted into another cup containing water, holding a broad bean cutting. To set up a new generation we transferred ∼20 asexual **L. fabarum** females into a fresh cup with ∼100 first-instar **A. fabae** and covered the cup with mesh (a cut section of tights) and a ventilated lid to prevent escape.

### 2.4. Experimental design

To test whether costs of facultative sexual reproduction constrain gene flow between asexual and sexual lineages, we quantified how lineages with evidence of sex performed across three generations, assessing their likelihood of reproductive failure and overall fecundity. We also evaluated whether antennomere number is associated with a history of facultative sex, and whether it can be used as a marker to detect hybridisation between sexual and asexual lineages.

Twenty-seven founding (F0) females were either mated with a male from the outbred sexual population (Fig. 2) or remained virgin. A subset of F1 females were genotyped to confirm facultative sex (this included multiple females from the same brood and F1 females used to found the F2). Individual F1 and F2 females were used to found subsequent broods and rearing was continued until F3. Offspring production was recorded across all generations to estimate fitness under facultative versus obligate asexual reproduction (previously inferred from mating status alone; Boulton 2025). Figure 2 provides an overview of the experimental design.

**Figure 2.**
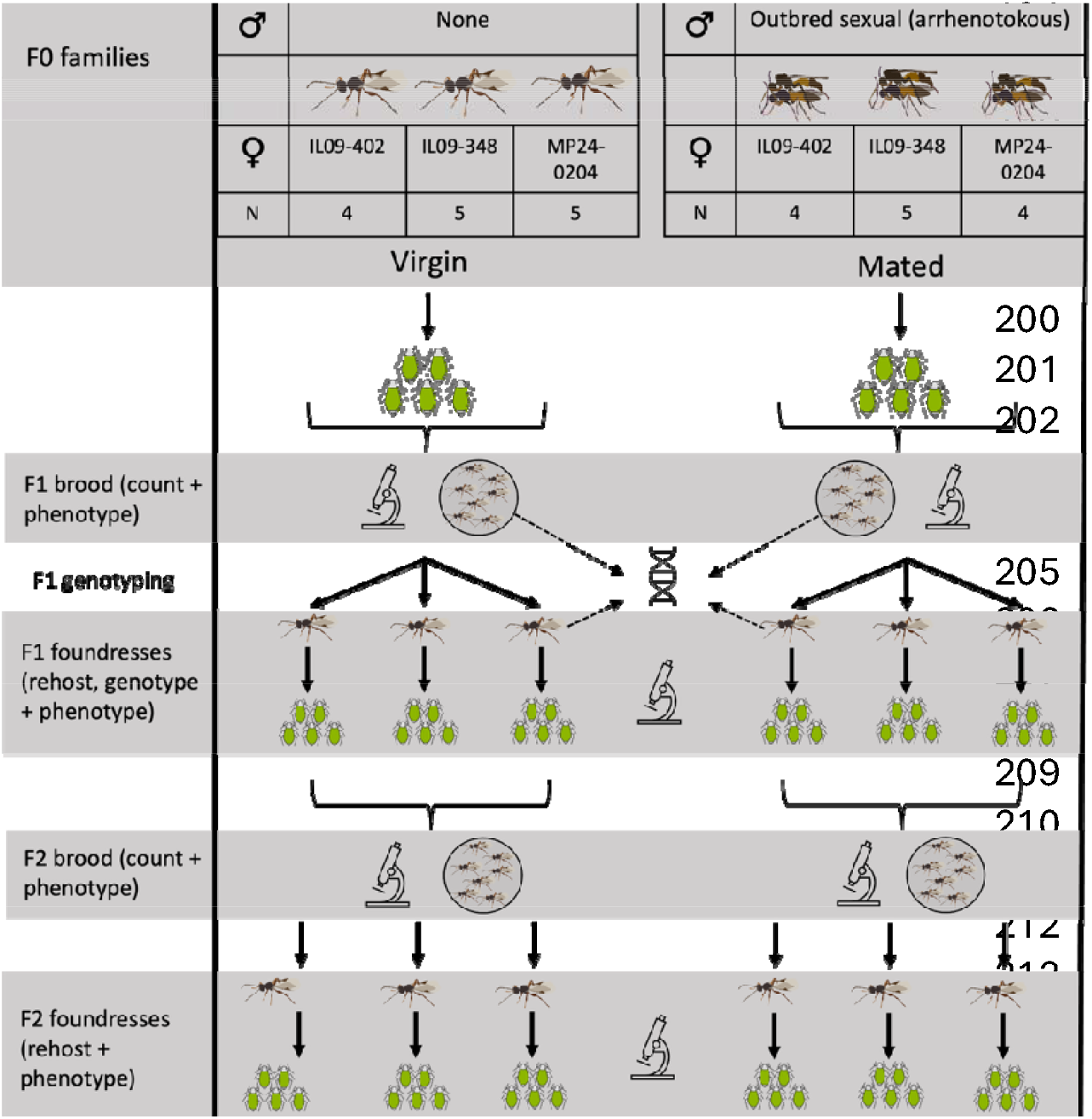
Schematic showing overall experimental design, number of generations and measurements taken each generation.

#### 2.4.1 F0 generation: setting up families

Across the three asexual lines (Table 1, Fig. 2), we established 27 F0 families, each founded by a single female. F0 females were either:

□ Virgin, or (N = 14)
□ Mated to a male from the outbred sexual (arrhenotokous) population (N = 13). Matings were carried out in clear cellulose pill capsules. F0 females and males were isolated as mummies and observed until emergence to ensure virgin status. All mating attempts were observed to confirm copulation.

Immediately after mating (or isolation for virgins), each F0 female was placed alone onto a fresh aphid colony in a culture cup (∼100 first instar *A. fabae*). After 2 weeks, resulting mummies were isolated to generate the F1 generation. Six F0 females produced no offspring (2 mated, 4 virgin), leaving 21 F0 families (11 mated, 10 virgin).

#### 2.4.2 F1 generation: establishing replicate lines

When F1 broods emerged, we established 59 replicate lines, taking 3–6 F1 females per family (all F1 females remained virgin). Each F1 founding female was placed alone into a fresh culture cup exactly as described for F0. Females oviposited alone for four days. We then removed and phenotyped the F1 founder females antennae and preserved their whole bodies in ethanol for genotyping (2.6). The remainder of the F1 brood was frozen, counted, and had their antennae phenotyped; additional F1 females (non-founders) were also genotyped.

#### 2.4.3 F2 and F3 generations

F2 founders were isolated as mummies, set up identically to F1 founders, and removed after four days; only antenna phenotype was recorded (no DNA extraction). All remaining F2 broods were counted and phenotyped. F3 was the final generation. All individuals were collected and phenotyped; no further propagation occurred (see Fig. 2).

### 2.5 Antennal phenotyping

We scored antennomere number under a compound microscope (model Leica M420 with Leica Apozoom 1:5 lens). For every female, both antennae (where intact) were examined and classified as sexual-12, asexual-13, or mixed following the scheme described in 2.2 (Fig. 1).

### 2.6 Genotyping

We genotyped 96 F1 females from 21 F0 families (30 of which founded F2 broods) at seven microsatellite loci using HotSHOT DNA extraction (Truett et al. 2000) and Qiagen Multiplex PCR. For each brood, between one and eight F1 females were genotyped, including both founders and their sisters, to determine whether offspring were produced exclusively asexually or via facultative sex, or whether broods contained a mixture of both reproductive modes.

Primer pairs were fluorescently labelled (forward) and unlabelled (reverse). Reactions used 1 µL DNA template in 11 µL reactions with Multiplex PCR Master Mix (Qiagen). PCR cycling conditions were as follows: 95°C 15 min → 30 cycles of 94°C 30 s, 56°C 90 s, 72°C 60 s → final extension 60°C 30 min.

PCR products were analysed by capillary electrophoresis at the MRC PPU sequencing facility (University of Dundee). Alleles were scored in Geneious™. *Lysi07* is closely linked to the thelytoky locus: asexual individuals carry a 184 bp allele, whereas sexual individuals carry a 192 bp allele Table 2). The presence of both alleles indicates that an individual was produced sexually (i.e., sperm use). Other loci provide secondary confirmation of sexual vs asexual origin (Table 2), as each asexual lineage has a characteristic multilocus genotype, deviations from this pattern, including the presence of unexpected or additional alleles, indicate genetic mixing consistent with facultative sex (see Table 2); *Lysi15* was uninformative in this dataset. Several individuals showed three alleles at polymorphic loci; preliminary flow cytometry suggests possible triploidy (K. Leung, unpublished).

**Table 2.** Microsatellite loci used in this study.

| Locus | Usefulness | Notes |
| --- | --- | --- |
| Lysi07 | Diagnostic | 184 = asexual allele; 192 = sexual allele; heterozygotes = facultative sex |
| Lysi06, Lysi08, Lysi13, Lysi16, Lysi03 | Supportive | Distinguish sexual vs asexual multilocus genotypes |
| Lysi15 | Uninformative | No differentiation between lineages |

### 2.7 Statistical analyses

All analyses were run in RStudio (Posit team, 2023) using the following packages: *nnet* (Venables & Ripley 2002), *brglm2* (Kosmidis 2025), *car* (Fox & Weisberg 2019), *lme4* (Bates et al. 2015), *glmmTMB* (Brooks et al. 2017) and *emmeans* (Lenth 2025), with model diagnostics conducted in DHARMa (Hartig 2024).

#### 2.7.1 Baseline analyses (stock populations)

We tested whether antennomere distributions differed across the four stock lines using bias-reduced multinomial regression using the package *brglm2* (dependent variable = antennomere category; predictor = line). Pairwise tests used multinomial Wald tests or Monte-Carlo r×c χ^2^ tests for lines exhibiting quasi-complete separation.

#### 2.7.2 F1 genotype–phenotype associations

We tested whether sexually vs asexually produced F1 females differed in antennal phenotype using 2×3 Monte-Carlo χ^2^ tests.

#### 2.7.3 Replicate-level models (F1–F3)

We ran a series of generalised linear mixed models (details in Table 3) where replicate (N = 59) was the unit of replication (family, N = 20, was included as a random effect). These models tested whether facultatively sexual replicate families were more likely to fail or experience low fecundity, or have a higher proportion females with sexual-12 type antennae. For models including F0 mating status, we additionally included line identity and its interaction with mating status to test whether the effects of mating varied across genetic backgrounds. Sample size constraints limited this analysis to F0 mating status, as comparable models including interactions for other predictors were not statistically appropriate due to small and unbalanced sample sizes.

**Table 3.**
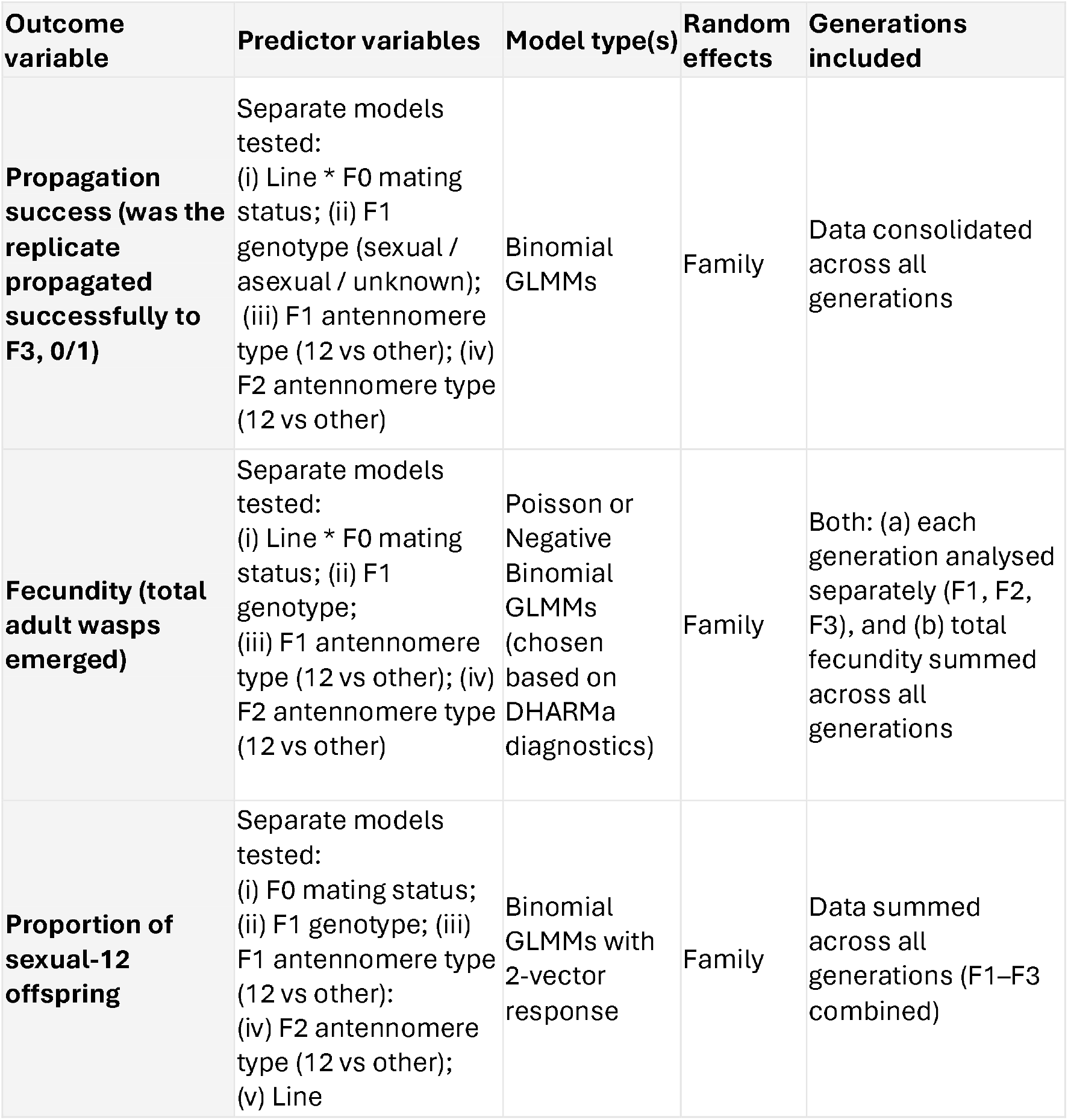
Summary of replicate-level statistical models.

#### 2.7.4 Family-level generational changes in antenna types

To test whether facultatively sexual lineages showed declining proportions of sexual-12 females over generations, we aggregated data at the level of family × generation. This was necessary because replicate lines were only defined from F2 onwards, following subdivision of F1 offspring into multiple replicate lineages; aggregation therefore provided a consistent unit of analysis across all generations.

We then used binomial GLMMs with family as a random effect to test for main and interaction effects of (i) F0 mating status × generation and (ii) evidence for sex in family × generation. Evidence of ‘sex in the family’ was defined as the presence of at least one F1 female within the family that was genotyped and confirmed to carry sexually derived alleles. These models test whether lineages with evidence of genetic mixing show changes in antennomere phenotype across generations, with declining proportions suggesting the costs of facultative sex limit the persistence of sexual alleles in asexual backgrounds.

## 3 Results

### 3.1 Stock population baselines

Across the four stock populations, the proportion of different antenna types varied across sexual and asexual lines (bias-reduced multinomial regression: LR χ^2^ = 143.98, df = 6, p < 0.0001). The sexual line consisted almost entirely of females with 12 antennomeres (sexual-12), whereas asexual lines produced mostly 13-antennomere females (asexual-13; Fig.□3A).

Coefficient estimates show that the sexual population contained few asexual-13 relative to sexual-12 (β = –4.40). In contrast, IL09-402 had much higher asexual-13 vs. sexual-12 (β = +2.52), consistent with the absence of sexual-12 females in this line. Because IL09-402 showed quasi-complete separation, we ran additional Monte-Carlo r×c χ^2^ tests for pairwise comparisons. These confirmed that the proportion of sexual-12 females in IL09-402 differed significantly from MP24-0204 (χ^2^ = 11.17, p = 0.003) but not from IL09-348 (χ^2^ = 5.30, p = 0.07). MP24-0204 and IL09-348 did not differ (χ^2^ = 3.17, p = 0.22).

### 3.2 F1 genotyping

We genotyped 96 F1 females from 21 families in total, 87 of which amplified successfully (44 from mated F0s, 43 from virgins). Of these, 18 F1 females from 7 families were produced via facultative sex and 69 females from 20 families via asexual thelytoky. As expected, most sexually produced females came from mated F0s (N = 16), although two sexually produced F1s originated from one virgin F0 female (this family was excluded from all further analyses). In only one family (derived from a mated F0 female) were all the genotyped F1 females produced sexually. In contrast, most mated F0 families showed a mixture of sexual and asexual F1s, and five produced no sexual offspring at all (Fig. 3B).

**Figure 3.**
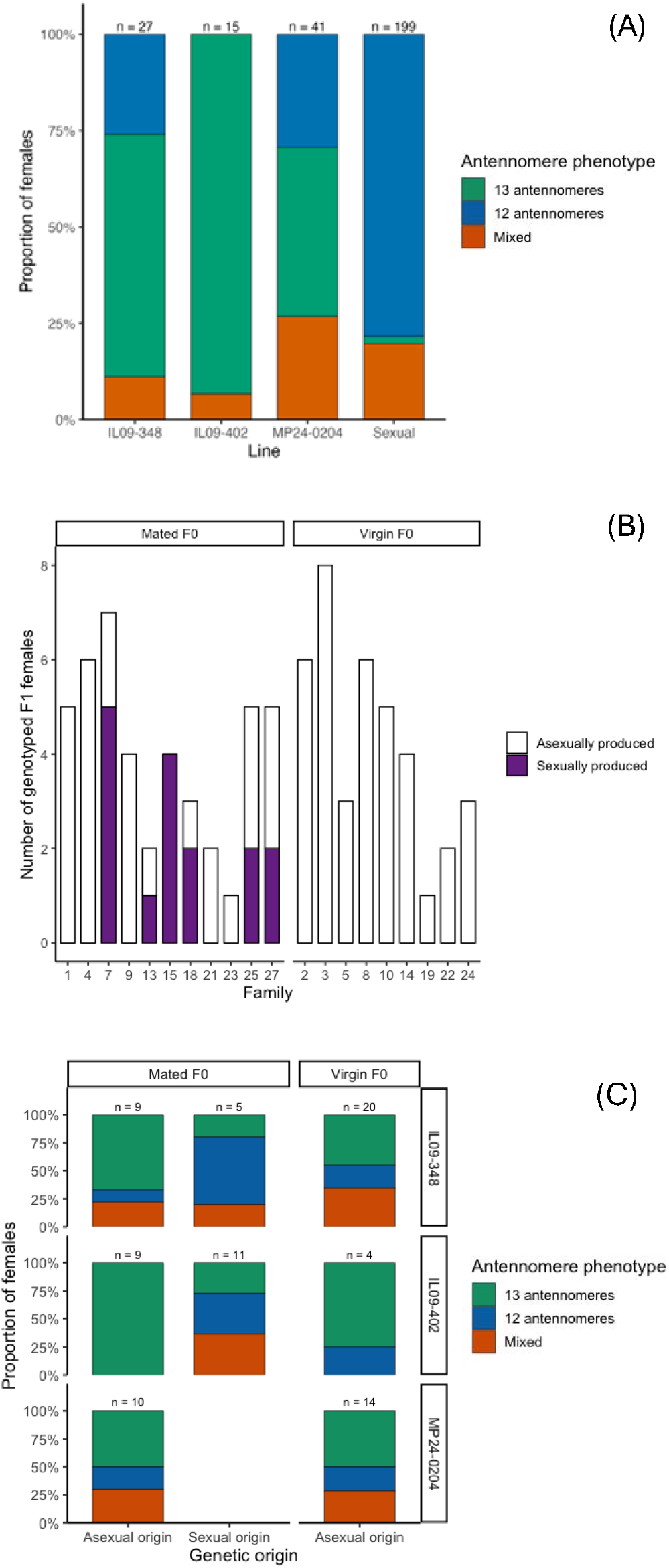
**(A)** Percentages of female wasps from four populations assigned to each antennomere phenotype category (sexual-12, asexual-13, or mixed). **(B)** Numbers of sexually and asexually produced genotyped F1 females from mated and virgin F0 females (by family). **(C)** Percentages of genotyped F1 females of sexual vs asexual origin from mated F0 and virgin F0 mothers assigned to each antenna phenotype. Sample sizes are displayed above each bar.

#### 3.2.1 Genotype-to-phenotype in F1

Antenna phenotypes of F1 females did not differ from the baseline distributions observed in the source asexual lineages in 3.1 (2×3 Monte-Carlo χ^2^ = 1.41, p = 0.50) but differed markedly from those observed in the sexual population (2×3 Monte-Carlo χ^2^ = 119.15, p = <0.0001). F1 phenotype distributions also differed from the expected midpoint between the sexual and asexual baseline distributions (i.e. the average of the sexual and asexual baseline phenotype frequencies; χ^2^ = 14.89, p < 0.001), indicating that overall F1 females more closely resembled asexual than sexual populations.

Although F1 females were overall very similar to asexual populations, antennomere phenotype was associated with genetic origin within the F1 generation (2×3 Monte-Carlo χ^2^ = 7.40, p = 0.02; Fig. 3C). Sexually produced F1 females were more likely to possess the sexual-associated 12-antennomere phenotype, whereas asexually produced F1 females were more likely to exhibit the asexual-associated 13-antennomere phenotype. Consistent with this pattern, sexually produced F1 females differed from both the asexual (χ^2^ = 6.28, p = 0.04) and sexual (χ^2^ = 24.36, p < 0.0005) baseline populations, but not from the expected midpoint between them (χ^2^ = 0.66, p = 0.81). In contrast, asexually produced F1 females did not differ from the asexual baseline population (χ^2^ = 1.38, p = 0.52), but differed from both the sexual baseline (χ^2^ = 130.00, p < 0.0001) and the midpoint expectation (χ^2^ = 17.78, p < 0.0001). F0 mating status itself was not associated with antennomere phenotype (χ^2^ = 0.41, p = 0.84).

#### 3.3.1 Reproductive failures and fecundity

Across all replicates, 25 (44%) failed to persist to F3, with failure rates of 37.5% in mated (12/32) and 52% in virgin (13/25) replicates. Reproductive failure was not significantly associated with F0 mating status, F1 genotype, or antennomere phenotype (all p > 0.17; Tables S1 & S2). Likewise, fecundity generally did not vary according to mating treatment or founder characteristics (Table S3).

In total we used 59 F1 females to found F2 replicate lines, we successfully genotyped 30 of these, 3 of which were produced sexually, 27 asexually. Although inference is limited by sample size, all three lineages founded by genotypically confirmed sexual F1 females failed before F3 and showed reduced cumulative offspring production relative to lineages founded by asexually produced F1 females (Table S3 & Fig. S1). This pattern is consistent with a fitness cost of sexual origin, but should be interpreted cautiously given the low number of confirmed sexual founders.

#### 3.3.2 Proportion of sexual-12 offspring

Across all generations combined, the proportion of sexual-12 offspring did not differ by F0 mated status (X^**2**^ = 0.29, df = 1, p = 0.59), F1 genotype (X^**2**^ = 0.41, df = 2, = 0.82) or F2 antenna phenotype (X^**2**^ = 1.72, df = 1, = 0.19). The only predictor with a significant effect was F1 antennomere phenotype: replicates founded by sexual-12 antennomere females had ∼90% higher odds of producing sexual-12 daughters (OR = 4.14, p = 0.04; Fig.□4).

**Figure 4.**
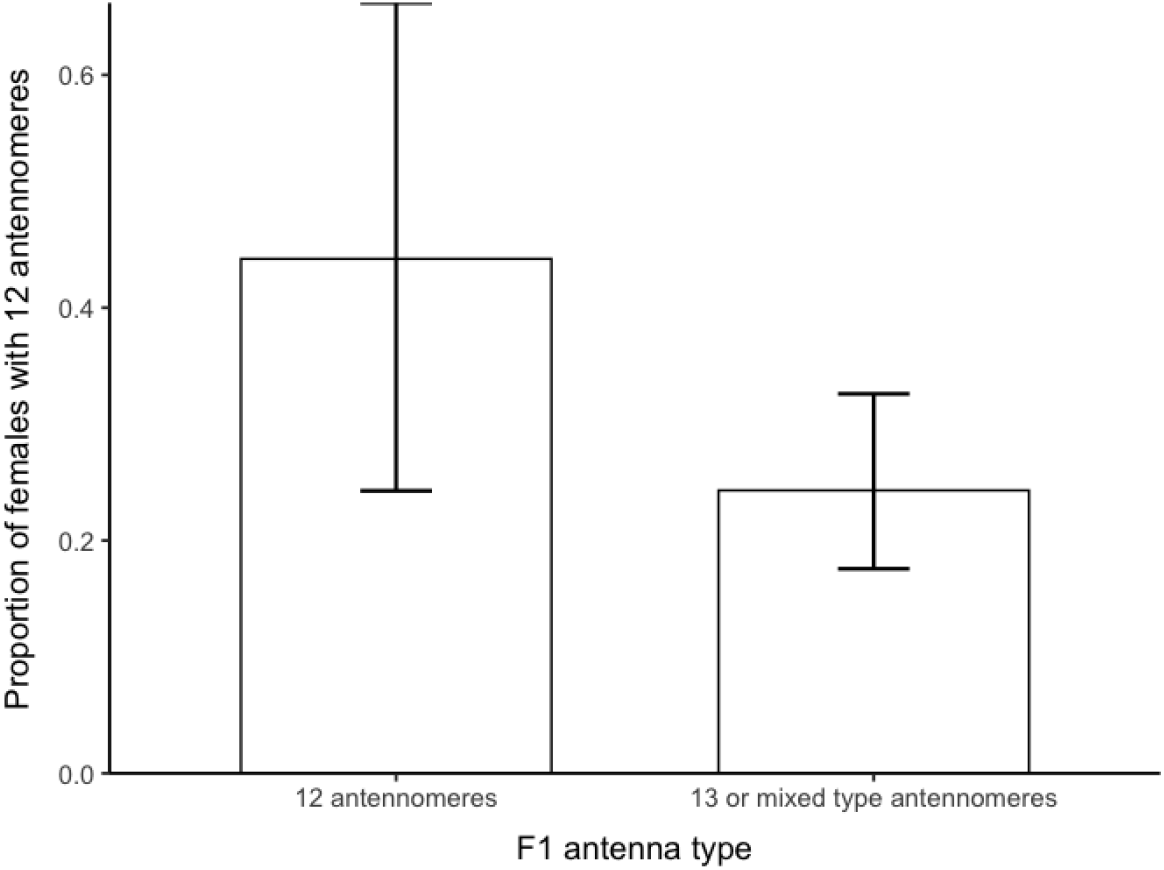
F1 females with sexual-12 antennomeres were ∼90% more likely to produce 11 antennomere offspring compared to females with asexual-13 or mixed type antennomeres. Proportions reflect estimated marginal probabilities from **emmeans** and error bars are confidence limits estimated by **emmeans**.

### 3.4 Family-level generational changes

To test whether sexually produced families lost the sexual-12 antennomere phenotype over generations, we analysed data summarised by family × generation. Neither F0 mated status or evidence for sex in the family (6 out of 20 families; Fig. 3B) had significant main effects. The only significant term was the interaction between sex in the family and generation (χ^2^ = 6.26, p = 0.04; Table□4).

**Table 4.** Results from binomial GLMMs testing for generational changes in the proportion of females with sexual-12 antenna according to founder mated status and whether there was evidence that any F1 females from the family were produced sexually.

| Model | Predictor | $X^2$ | df | p |
| --- | --- | --- | --- | --- |
| F0 mated status | F0 mated status | 1.70 | 1 | 0.19 |
|  | Generation | 2.46 | 2 | 0.29 |
|  | F0 mated status * generation | 5.65 | 2 | 0.06 |
| Sex in family | Sex in family | 0.55 | 1 | 0.46 |
|  | Generation | 2.45 | 2 | 0.29 |
|  | **Sex in family \***<br>**generation** | **6.26** | **2** | **0.04** |

Pairwise contrasts show no significant differences in the proportion of females with sexual-12 type antenna between families with and without evidence of sexual reproduction within individual generations (all p > 0.12), but patterns across generations did differ between the two groups. In families with evidence of sex, the proportion of females with sexual-12 type antenna declined significantly from F1 to F2 (odds ratio = 3.13, p = 0.026), whereas no such change was observed in families without evidence of sex (Fig. 5).

**Figure 5.**
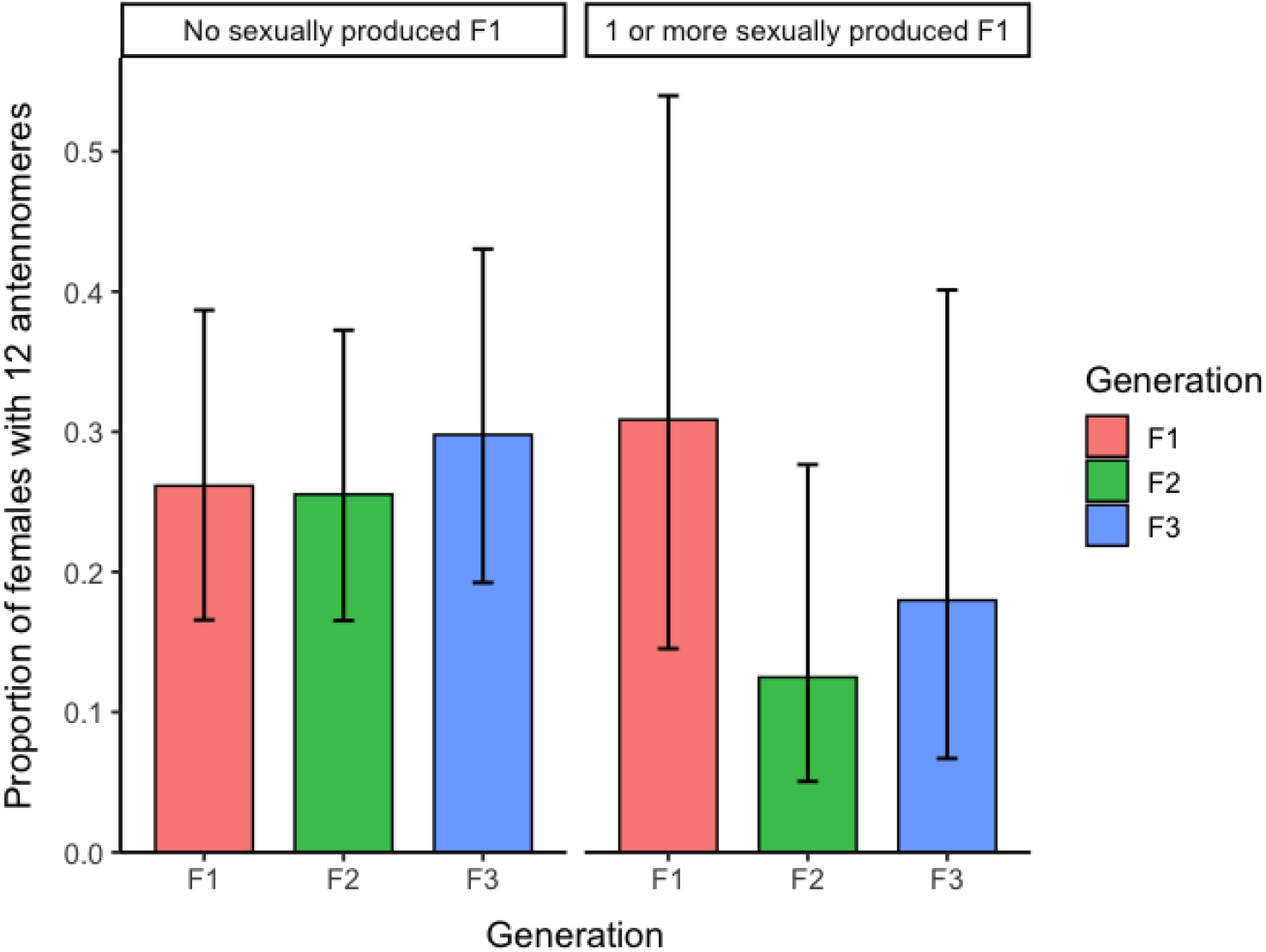
Proportion of females with 12 antennomeres across generations for families with no genotypic evidence of sexual reproduction versus families in which at least one F1 female was genotyped as sexually produced. Proportions are estimated marginal probabilities from *emmeans*, with error bars representing confidence intervals.

### 4.0 Discussion

### 4.1 Summary

Facultative sex in *Lysiphlebus fabarum* creates an opportunity for gene flow between sexual and asexual lineages and may be useful for answering questions about how often facultative sex occurs, whether and under what conditions sexual alleles can become stably incorporated into asexual genomic backgrounds, and how cryptic facultative sex can be detected in wild populations (Neiman et al. 2014; Kokko 2020; Freitas et al. 2023; Wilner et al. 2025; Piezko et al. 2025). To begin addressing these questions requires an understanding of the frequency and consequences of facultative sex in *L. fabarum*. As such, the aims of this study were fourfold, we wanted to determine (1) how often mating results in facultative sex, (2) whether all offspring of mated thelytokous females are produced sexually, (3) and whether fitness costs are associated with **the genetic consequences of** facultative sex itself (rather than simply with having a mated mother). Finally, we aimed to test whether (4) antennomere number could be used to track sexual alleles through predominantly asexual populations, with the prediction that fitness costs and constraints on facultative sex (i.e. 1-3) would result in declining frequencies of the sexually-associated 12-antennomere phenotype across generations.

By genotyping individual F1 females, we found that (1) mating did not always result in **sexually produced offspring**, with sexual reproduction confirmed in only seven of eleven families produced by mated F0 females. Moreover, even when **a female produced offspring sexually**, (2) broods typically contained a mixture of sexual and asexual daughters. Facultative sex also carried fitness costs (3). While sample sizes were low, the 3 lineages founded by sexually produced F1 females exhibited significantly lower fecundity and none of these lines persisted for all three generations. An important limitation is that we used only a single sexual population, meaning that the costs observed here cannot be disentangled from potential outbreeding effects. Consequently, our results demonstrate a cost of producing mixed sexual-asexual genotypes, but do not allow us to determine whether such costs are a general consequence of facultative sex or reflect incompatibilities between the lineages used in this study.

In terms of (4), antennomere number was associated with reproductive mode, with sexually produced F1 females more likely to possess the sexually-associated sexual-12 antennomere phenotype. The mixed-antenna phenotype showed no consistent associations with reproductive mode or inheritance, implying that it may reflect developmental variability rather than a stable or heritable state (Minelli 2017). Additionally, F1 females with sexual-12 antennae produced higher proportions of sexual-12 offspring, suggesting that this trait is heritable. Our prediction that the frequency of the sexual-12 antennomere phenotype would decline over time after facultative sex (due to fitness costs/constraints; 1-3) was supported. Mated families where at least one F1 female had a mixed genotype showed a significant decline from around 30% sexual-12 females in the F1 generation to approximately 12% in the F2 generation, followed by an increase to around 18% in the F3 generation. Although the trajectory was as predicted, the frequencies of sexual-12 antennomere females were generally as high or higher in virgin families (∼25–30% across all three generations). Whilst the occurrence of sexual-12 antennomere females at reasonable frequencies in confirmed asexual lines limits the conclusions that can be drawn about the persistence of sexual alleles from antennomere number alone, the observed pattern is consistent with the idea that facultative sex produces low fitness genotypes (Otto & Lenormand 2002; Otto 2021; Parée & Teotónio 2025) resulting in declining frequencies of hybrid phenotypes.

One possibility is that F2 hybrid offspring have low viability and are less likely to survive to eclosion. This is consistent with the reduced fecundity and persistence of facultatively sexual lineages observed in this study and could also explain why the frequency of sexual-12 females falls below that seen in equivalent families with no evidence of facultative sex. Sexual-12 females occur at a background frequency of approximately 25% in both virgin families and stock populations, indicating that this phenotype is not exclusively associated with hybrid ancestry. Consequently, in families that experienced facultative sex, the pool of sexual-12 females may comprise both background sexual-12 individuals and individuals carrying sexually derived alleles. If the latter group experiences reduced viability, selection would disproportionately remove hybrid-derived sexual-12 females while leaving the background component largely unchanged. This would generate a temporary decline in the overall frequency of sexual-12 females, consistent with the reduction observed in F2.

### 4.2 Costs and constraints of facultative sex

In earlier work we showed that facultative sex in *L. fabarum* can be costly, as putative F1 hybrids (they were not genotyped, their mothers had mated) were more likely to fail to reproduce (Boulton 2025). In the current study our results add nuance as we were able to genotype a proportion of individual F1 mothers. Although only 3 of the genotyped F1 founders were confirmed to have been produced sexually, none of these were successfully propagated for all 3 generations and fecundity was significantly lower in these lines compared to those founded by asexually produced F1 females. These results add to a growing body of evidence suggesting that facultative sex is not the “best of both worlds,” but instead introduces novel fitness costs not seen under obligate sexual reproduction (Lehtonen et al. 2012; Hartfield 2016; Parée & Teotónio 2025).

An additional novel finding of this study was that facultative sex in *L. fabarum* is genuinely facultative: not all mated females produce hybrids, and of those that do, most produced broods with mixed genotypes. In our analyses among mated F0 families, five produced no hybrid F1 daughters, five produced mixed broods, and in only one brood were all genotyped females hybrids. These results reinforce the idea that thelytokous females can produce both sexual and asexual daughters even in the same brood and that the allocation of sexual eggs is highly variable. Mechanistically, this is consistent with the ancestral arrhenotokous sex-allocation system where unfertilised eggs become haploid males and fertilised eggs diploid daughters (de la Filia et al. 2015); under thelytoky this results in unfertilised eggs becoming diploid asexual daughters and fertilised eggs hybrid daughters.

The facultative production of mixed broods has wider evolutionary consequences. Under facultative thelytoky, the ancestral fertilisation system allows females to hedge their bets by producing some sexually derived offspring even when the fitness of those hybrids is uncertain. This aligns with bet-hedging arguments for the benefits of sex, in which producing offspring with diverse genotypes increases the geometric mean fitness across generations under changing conditions (Bell 1982; Williams 1975), while asexuals maintain a high arithmetic mean fitness but could fail to reproduce entirely if conditions change. In *L. fabarum* however, a single female can effectively do both, reaping the short-term efficiency of parthenogenesis while still generating occasional genetic novelty through sex.

However, this flexibility comes with costs. As we saw in this study, hybrid daughters typically have lower fecundity and may even be sterile. These costs are likely to become worse when they are in direct competition with their asexually produced sisters, particularly under natural conditions where hosts are patchy, seasonal, and frequently limiting (Jervis et al. 2023). In our experiment, lineages were maintained as isofemale lines and thus did not experience direct competition, meaning these effects are likely to be conservative relative to natural conditions. In nature, such competitive asymmetries are likely to reduce the persistence of sexual alleles introduced into asexual genomic backgrounds, even when facultative sex does occur. As such, even if facultative sex can generate hybrids capable of reproducing, there are many factors working against stable incorporation of sexual alleles. Taken together, these constraints, fitness variation among hybrids, variable sex allocation, and ecological competition, may help explain why facultative sex is rarely detected in natural populations.

### 4.3 Conclusions and future directions

Overall, our study demonstrates that facultative sex in *L. fabarum* does occur, can generate hybrids and may lead to some introgression of sexual alleles into asexual lineages. However, the process is inconsistent, often costly, and easily disrupted. Antenna phenotype is not ideal for tracking introgression in this system, but is a useful preliminary indicator when combined with genotyping.

An important next step will be to maintain confirmed facultatively sexual lines across multiple generations and track allele transmission using higher-resolution genomic approaches. Genomic data for entire broods would also provide a more complete picture of how sexual and asexual offspring are allocated within families and how introgressed alleles persist over time. A complementary approach would be to identify or establish a “clean” asexual-13 lineage, allowing the effects of facultative sex on sexual-associated phenotypes to be tested more rigorously.

Equally important is identifying the mechanistic basis of the fitness costs associated with facultative sex. Preliminary work suggests that triploidy may occasionally occur in this system (pers. comm. Kelley Leung), while outbreeding depression, genetic incompatibilities, and other cytogenetic irregularities are also plausible (Lynch 1991; Darras et al. 2023). Distinguishing between these possibilities is important because they may result in different evolutionary consequences. Cytogenetic abnormalities such as triploidy might impose severe and largely irreversible fitness reductions or complete reproductive failure, whereas incompatibilities arising through outbreeding depression may depend on the degree of divergence between lineages and could be less pronounced among closely related populations. We were unable to directly test the costs of outbreeding in sexuals because only a single outbred sexual population was available. Determining whether outbreeding similarly reduces fecundity by crossing different sexual lineages would help clarify the extent to which outbreeding depression contributes to the fitness costs observed under facultative sex. Finally, field sampling will be essential to determine how frequently facultative sex occurs under natural conditions and whether hybrid lineages exist and persist outside of the laboratory.

