## supplementary figure S1 for "Mating allows asexual female wasps to produce offspring both sexually and asexually, but their sexual daughters have reduced fecundity"

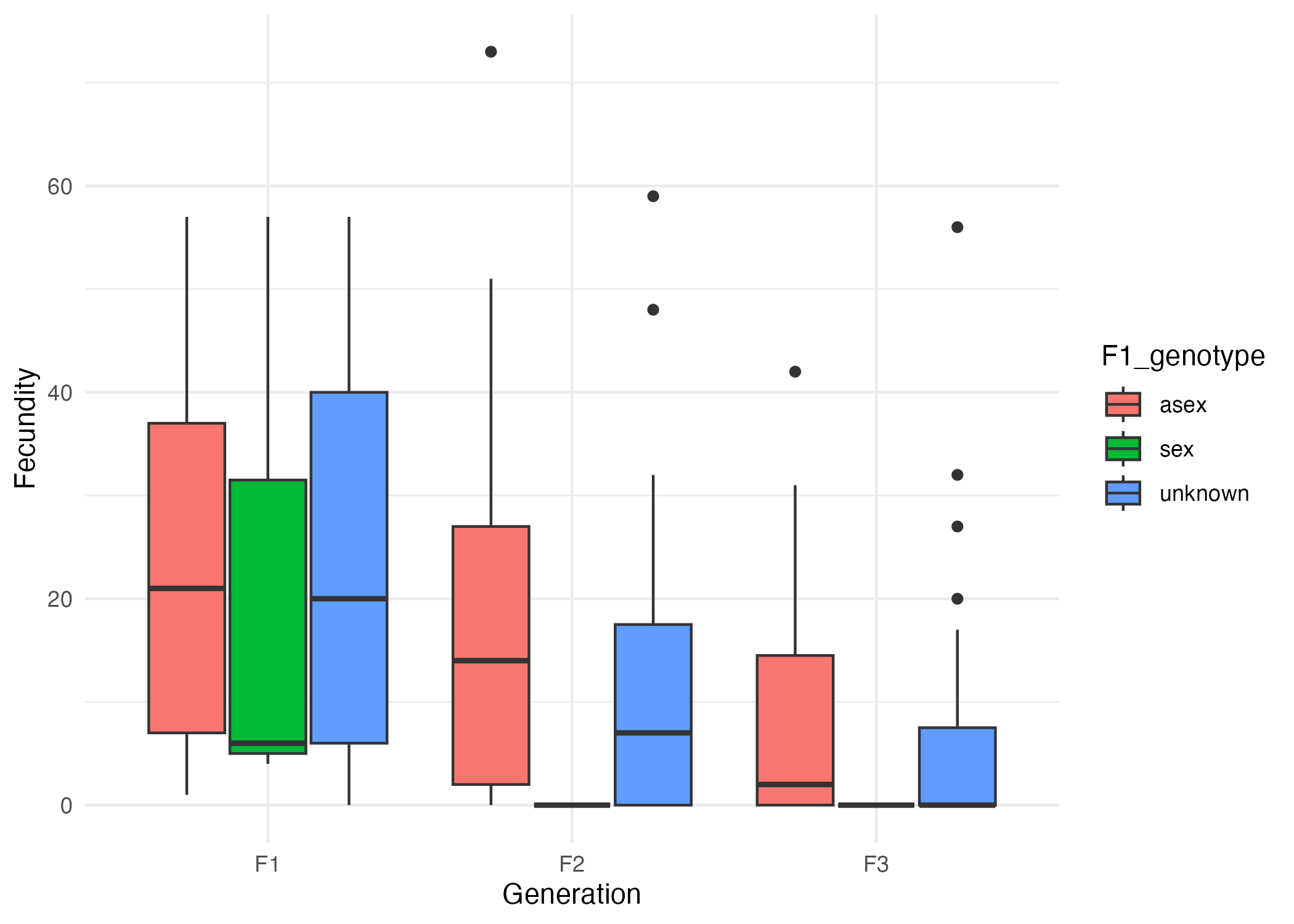

**Supplementary figure S2 |** Offspring production (fecundity) according to F1 female genetic status (produced sexually, asexually or unknown) across 3 generations. Bars show estimated marginal means with 95% CIs from *emmeans*.
