## supplementary table S1 for "Mating allows asexual female wasps to produce offspring both sexually and asexually, but their sexual daughters have reduced fecundity"

**Supplementary table S1 |** Replicate lines based on founding females characteristics and whether or not they were successfully propagated for all 3 generations

| Variable | Level | Successes | Failures | % sucess |
| --- | --- | --- | --- | --- |
| F0 mated status | Virgin | 11 | 14 | 56% |
|  | Mated | 15 | 17 | 53% |
| F1 genotype | Sexual | 0 | 3 | 100% |
|  | Asexual | 16 | 15 | 48% |
|  | Unknown | 10 | 13 | 57% |
| F1 antenna | Sexual_12 | 2 | 1 | 33% |
|  | Asexual_13 | 17 | 14 | 45% |
|  | Mixed | 3 | 3 | 50% |
|  | Unknown | 4 | 13 | 77% |
| F2 antenna | Sexual_12 | 4 | 2 | 22% |
|  | Asexual_13 | 13 | 5 | 28% |
|  | Mixed | 3 | 2 | 40% |
|  | Unknown | 6 | 22 | 79% |
