## supplementary table S2 for "Mating allows asexual female wasps to produce offspring both sexually and asexually, but their sexual daughters have reduced fecundity"

**Supplementary table S2 |** Estimated marginal means, odds ratios and test statistics for pairwise contrasts for fecundity differences across lines extracted from GLMMs using *emmeans*

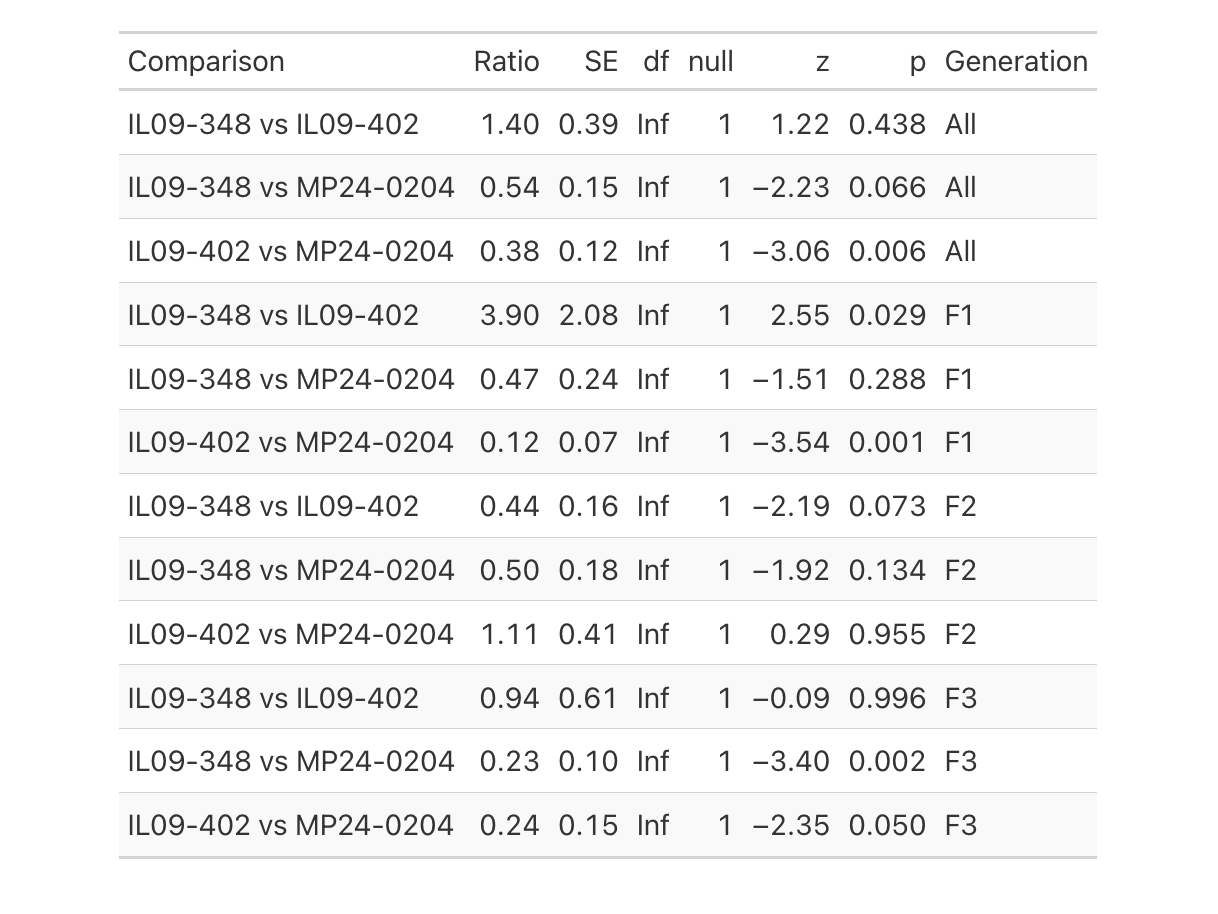
