## supplementary table S3 for "Mating allows asexual female wasps to produce offspring both sexually and asexually, but their sexual daughters have reduced fecundity"

**Supplementary table S3 |** Estimated marginal means, odds ratios and test statistics for pairwise contrasts for fecundity differences across lines and generations extracted from GLMMs using emmeans

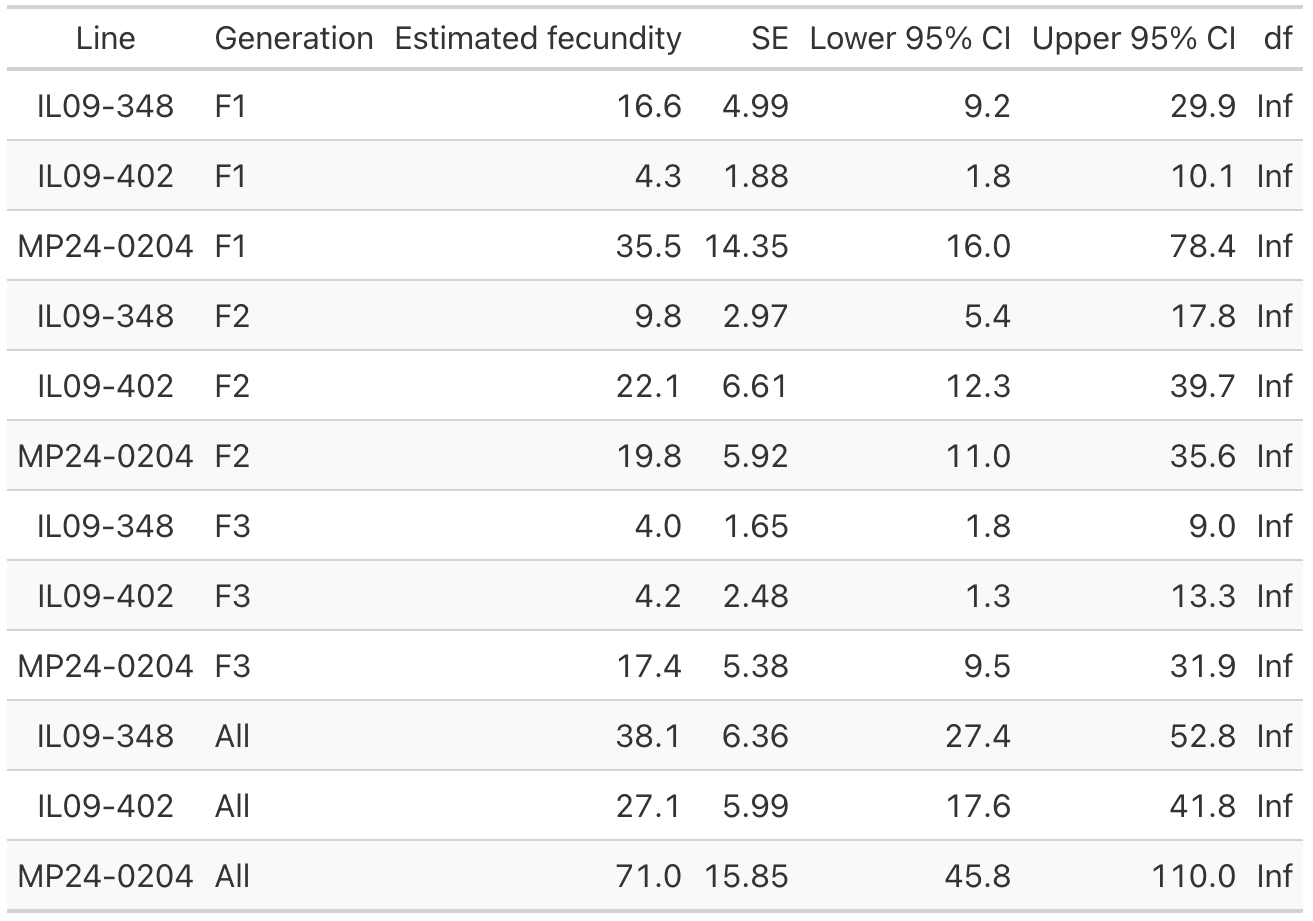
